# A Coupled Network of Growth Transform Neurons for Spike-Encoded Auditory Feature Extraction

**DOI:** 10.1101/303453

**Authors:** Ahana Gangopadhyay, Kenji Aono, Darshit Mehta, Shantanu Chakrabartty

## Abstract

This paper builds upon our previously reported growth transform based optimization framework to present a novel spiking neuron model and demonstrate its application for spike-based auditory signal processing. Unlike conventional neuromorphic approaches, the proposed Growth Transform (GT) neuron model is tightly coupled to a system objective function, which results in network dynamics that are always stable and interpretable; and the process of spike generation and population dynamics is the result of minimizing an energy functional. We then extend the model to include axonal propagation delays in a manner that the optimized solution of the system or network objective function remains unaffected. The paper characterizes the model for different types of stimuli, and explores how changing different aspects of the cost function can reproduce known single neuron dynamics. We then investigate the properties of a coupled GT neural network that can generate spike-encoded auditory features corresponding to the output of a gammatone filterbank. We show that the discriminatory information is not only encoded in the traditional spike-rates and interspike-interval statistics, but is also encoded in the subthreshold response of GT neurons for inputs that are not strong enough to elicit spikes. We also demonstrate that while different forms of coupling between the neurons could produce compact and energy-efficient representation of the auditory features, the classification performance for a speaker recognition task remains invariant across different types of coupling. Thus, we believe that the proposed GT neuron model provides a flexible neuromorphic framework to systematically design large-scale spiking neural networks with stable and interpretable dynamics.

## I. INTRODUCTION

One of the main paradigms in the design of neuromorphic systems is to be able to achieve robust and efficient recognition performance by mimicking the architecture and the dynamics of biological neural networks. For example, neuromorphic implementations of the cochlea ([1]–[5]) closely mimic the neural dynamics observed in the auditory pathway where sound waves set up mechanical vibrations that are converted to electrical impulses by hair cells present along the cochlear basilar membrane in the inner ear ([6]). Specifically, continuous-time filterbanks have been used to emulate the tonotopic organization observed in the human auditory pathways, where groups of neurons in close proximity respond preferentially to a specific range in the frequency spectrum ([7]). This is illustrated in Figure 1A by an array of spiking neurons that processes the outputs of the filterbanks to produce neural impulses that encode the auditory features. In biological systems, the neural impulses are carried by auditory nerves through several auditory centers before reaching the higher processing stages in the cerebral cortex, undergoing several stages of complex transformations that are important for perception, identification, localization and segregation of sound. In synthetic auditory systems, as shown in Figure 1A, specific higher processing functions like speech or speaker recognition are emulated using standard machine learning modules like support vector machines (SVM) or convolutional neural networks (CNN). Irrespective of the type of the back-end classifier, the system recognition performance is ultimately determined by the discriminatory information present in the auditory features that are encoded by the spiking neuron models. In literature, neuronal models of varying degrees of complexity, ranging from the classic Hodgkin-Huxley model ([8]), FitzHugh-Nagumo model ([9]) and Izhikevich models ([10]) to the simple integrate-and-fire model ([11]) have been used. However, practical implementations of the cochlea usually resort to a simpler form of these models ([12]), where only the limit-cycle statistics like average rates or inter-spike intervals of the spike-trains produced are then used for generating auditory features for recognition or classification([13], [14]). At this level of abstraction, it is not evident how the shape, the nature and the dynamics of each individual spike is related to the overall system objective, and how a population of neurons when coupled together can self-optimize itself to produce an emergent spiking or population response ([15]). This is important for audition since it has been reported that each stage of the auditory pathway also exhibits limit cycles and subthreshold oscillations, both of which are believed to be important for precise encoding of acoustic information ([16], [17]). How these synaptic connections and the rich repertoire of dynamics encode speech signals for discrimination is a problem that remains unsolved, as is how to interpret and control such neural responses in a manner that can achieve a desired system objective.

**Figure 1.**
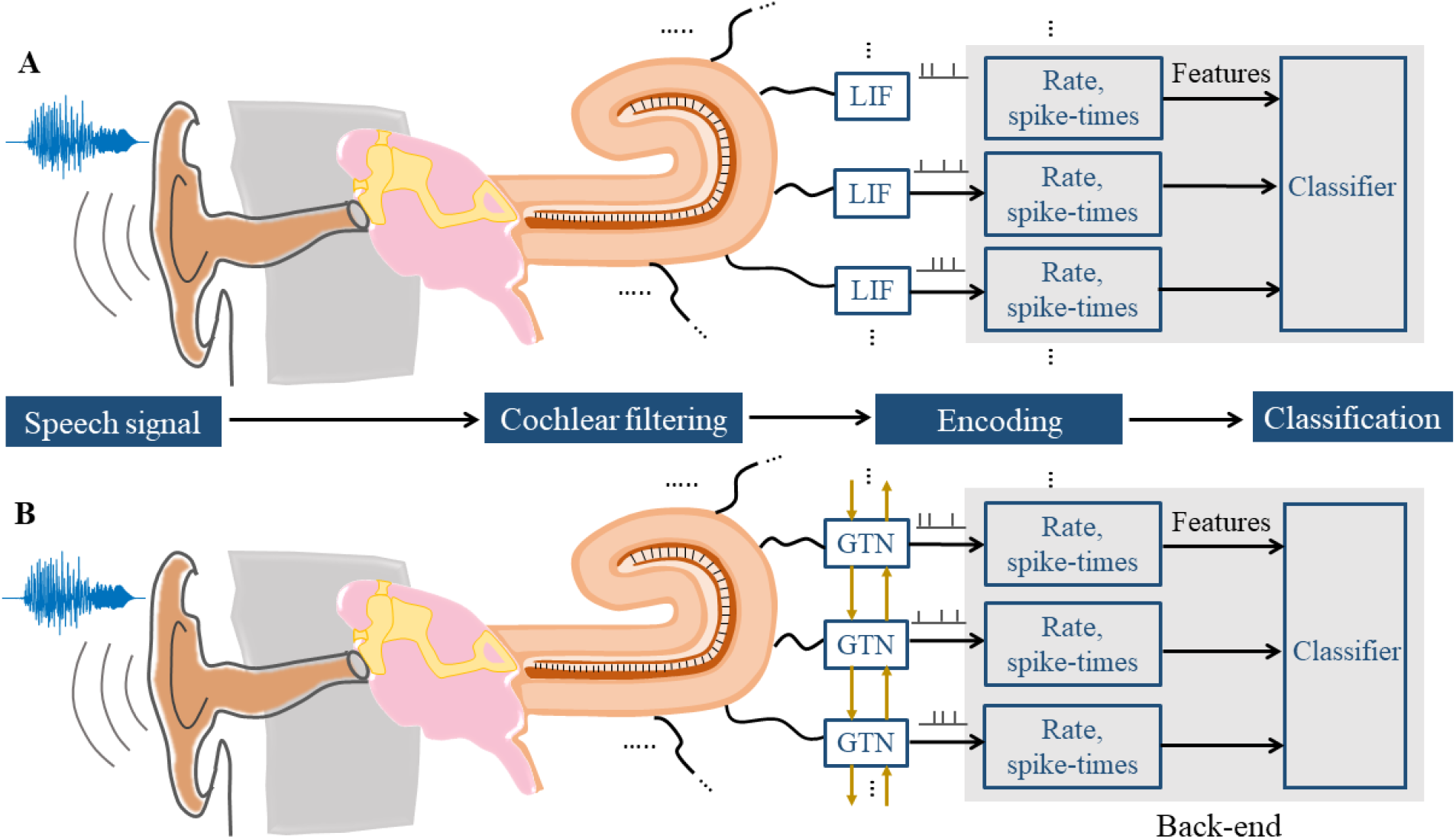
(A) Standard neuromorphic auditory feature extraction using leaky integrate-and-fire (LIF) neuron models followed by a DCT-based decorrelation and signal classification. (B) Proposed architecture using a coupled network of Growth Transform (GT) neurons to achieve a more energy-efficient auditory encoding.

In this paper, we approach the process of spike generation and encoding in a neuron model from a perspective that is fundamentally different compared to traditional dynamical systems formulations - by designing an energy functional for a network of neurons which produces an emergent spiking dynamics when minimized under realistic physical constraints. This is motivated by the fact that physical processes occurring in nature have an universal tendency to move towards a minimum-energy state - defined by the values taken by the process variables - that is a function of the inputs and the state of the process. Energy-based frameworks are the backbone of most machine learning models, which perform inference by minimizing an energy functional that capture dependencies between the variables ([18]). Energy consideration has either been largely overlooked in conventional representations of spiking neural networks, or been modeled as an external resource to constrain and control neural activity ([19]). This is despite the fact that neural signaling is metabolically expensive, with the neuron expending a significant fraction of its net energy utilization in the activation/deactivation of ion channels for spike generation and transmission ([20]). It therefore makes sense to study the process of spike generation and neural representation of excitation in the context of a network-level energy-minimization problem. ([21]) takes a similar approach by considering an energy function of spike-based network states, where individual neuron states are assumed to be binary in nature (spiking or not spiking). However, there is a lack of an unified model in literature that treats the spike generation process as the direct derivative of an energy functional of continuous-valued neural variables (e.g. membrane potential), that can moreover be used to mimic known neural dynamics from literature.

In ([22]), we proposed a model of a growth transform (GT) neuron whose dynamical and spiking responses were shown to be strongly coupled with the network objective function. It was shown that while each individual neuron traverses a trajectory in the dual optimization space, the overall network traverses a trajectory in an equivalent primal optimization space. As a result, the network of growth transform neurons was shown to solve classification tasks, while producing stable, unique but interpretable neural dynamics like noise-shaping, spiking and bursting. In this paper, we first introduce a generic energy (cost) function for spiking neurons by applying a current balance equation at each neural compartment in the network, that stochastically maintains a charge-equilibrium in the cells. We then extend the model to incorporate axonal delays without affecting the network stability by exploiting the properties of growth transforms. We show that the GT neural network can exhibit known response characteristics similar to biological neurons by altering different aspects of the energy functional. Finally, as illustrated in Figure 1B, we explore a coupled network of GT neurons for generation and encoding of auditory features that can be used for discriminating speakers. We show that as the input stimulus varies over time, the spiking pattern of the growth transform neuron adapts, effectively tracking and transforming the input, while always being constrained by a system-level objective. Apart from an uncoupled network where the neurons track individual filterbank outputs, we show results for the speaker recognition task with coupled networks having sparse random excitatory and/or inhibitory connections between neurons. This formulation enables us to explore different patterns of connectivity between neurons, which may lead to a better or more compact stimulus representation. For example, inhibitory connections reduce the overall firing rate of the network, but produces similar classification performance as the uncoupled network.

Another novel aspect of this work compared to other neuromorphic approaches is the use of subthreshold response generated by GT neurons for auditory discrimination in addition to spike-based statistics. When the stimulus is not strong enough to elicit spiking from a neuron, its subthreshold activity still encodes information about the stimulus that cannot be obtained from its spiking activity only. Electrophysiological recordings across different sensory modalities and brain regions have shown that subthreshold response of a neuron is stimulus-specific and highly robust [23]–[25]. Rate encoding is a special case of this feature where the membrane potential trace is thresholded to select only the spikes. In this paper, we show that while information carried by the rate or timing of spikes can indeed encode useful information from speech signals, inclusion of subthreshold response results in a finer discrimination between speech signals from different speakers when compared to rate encoding.

## II. METHODS

In this section, we introduce an energy functional for the GT neural network and then derive the basic model of a GT neuron as a result of its minimization. We then extend this model to include axonal propagation delays by exploiting properties of growth transforms.

### A. Energy function for Growth Transform spiking neuron model

The theory underlying growth transforms and its connections with different forms of system objective functions, network dynamics (spiking, bursting and noise-shaping) have been discussed in detail in [22]. Here we summarize the key concepts underlying the derivation of the GT neuron model based on the energy minimization framework. We will abstract the physics of neuronal spike generation and spike propagation using a multi-compartmental model as shown in Figure 2. Here each compartment could model a neuron which reacts to external stimuli and propagate signals to other compartments (neurons) by producing a rapid change in their transmembrane electrical potential difference (known as an action potential or a spike). Each compartment in the model comprises of ion channels, as shown in Fig. 2, whose roles are to equilibrate the charge across the membrane to its resting state. Also, in this simplified model we assume that the coupling between the compartments *i* and *j* is given by a transimpedance variable *Q_ij_*.

**Figure 2.**
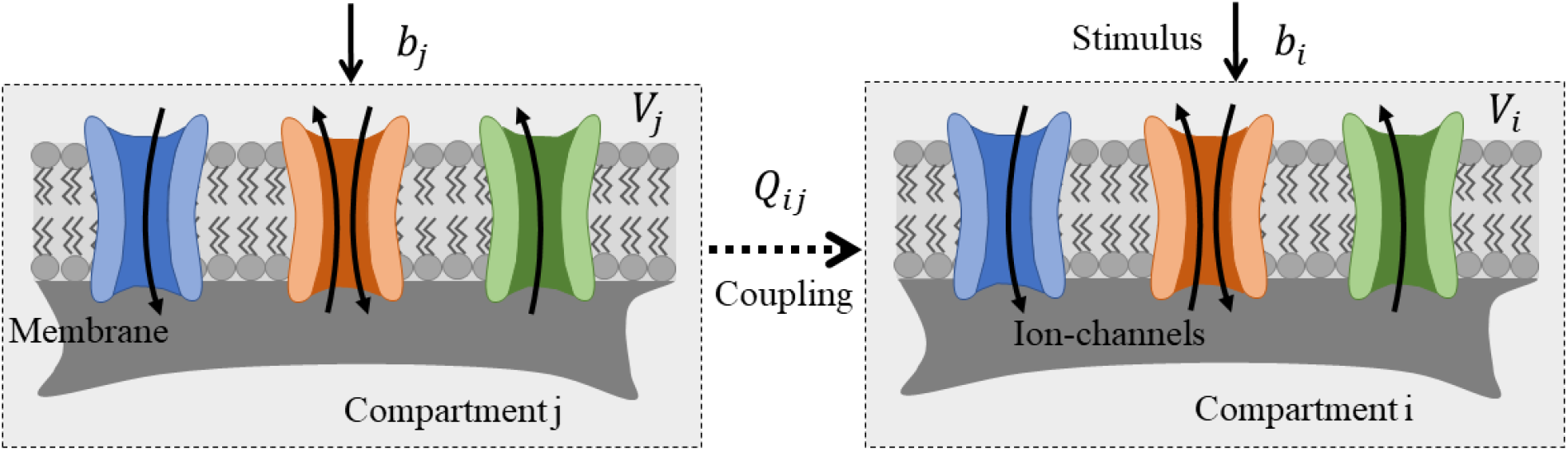
A subsection of the compartmental network is shown. Each neuronal compartment receives its own external stimulus *b*. The j-th compartment is coupled to the i-th by a coupling strength *Q_ij_*.

Assuming that the charge across the membrane remains stable, each compartment maintains a current-balance where the sum of the time-average of the net transmembrane current due to ion-channel activity and the time-average of the time-varying input stimulus received by a compartment (neuron) is zero. Defining the time-averaging operator *ε*(·) as an expectation of a signal *X(t)* computed over a time range [0 *T*] as

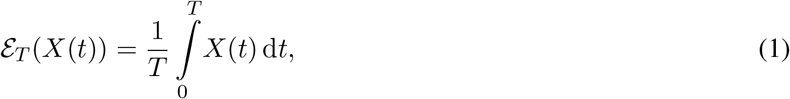

the current-balance equation for the *i*-th neuron can be expressed in terms of its instantaneous membrane potential *V_i_(t)* as

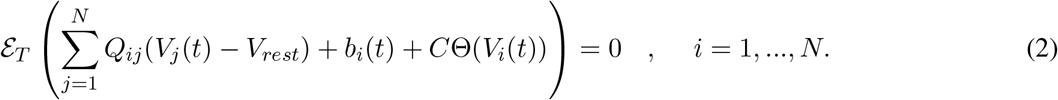

Here, *N* is the total number of neurons in the network, 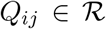 denotes the contribution of the pre-synaptic potential at neuron *j* to the net excitation received by the post-synaptic neuron 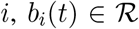 is the instantaneous external current stimulus, and Θ(·) refers to the net transmembrane ionic current flow that maps the instantaneous membrane potential *V_i_(t)* into a spike response. *V_rest_* denotes the resting membrane potential. Note that we are abstracting the effect of multiple ion-channels (for example sodium, potassim or calcium) into a single ion-channel current function. An illustration of the compartmental model is given in Figure 2. *C* is a tunable parameter that, as we will show later, models the excitability of a compartment (neuron) to its input. We additionally consider the membrane potential to be always bounded as follows

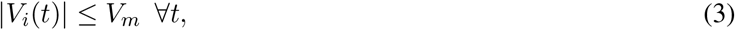

for some constant *V_m_* > 0. The principle is similar to the neuron membrane potential being bounded by the ionic potentials on either side of the cell membrane that mediate the changes in membrane potential. For the sake of simplicity, we consider the membrane potential to be normalized, such that

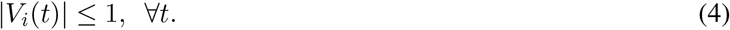

We can decompose the response *V_i_(t)* of the *i*-th neuron and the bias term *b_i_(t)* into two differential components as follows

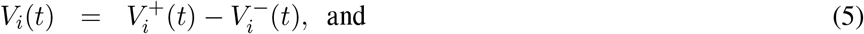

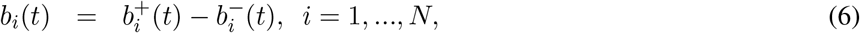

where 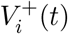 and 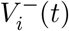 satisfy the following constraints:

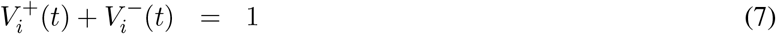

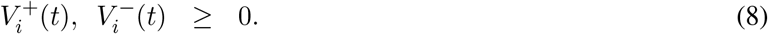

Note that the Eqs 7 and 8 ensure that the decomposition of *V_i_(t)* is unique, whereas the decomposition of *b_i_(t)* is non-unique and does not affect the solution. Similarly, the resting membrane potential (also normalized w.r.t. *V_m_*) can be uniquely decomposed as 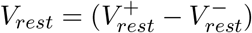. Eq. 2 can thus be rewritten as

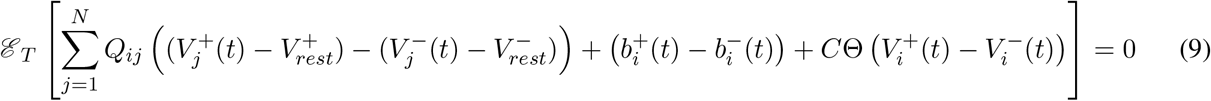

For simplicity, we choose a piece-wise linear form of Θ(·) such that it can be decomposed in the following way

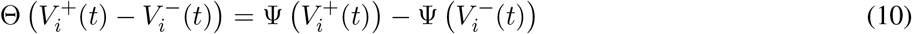

for some function Ψ(·). Thus, solving Eq. 9 is equivalent to satisfying the following (stronger) set of conditions:

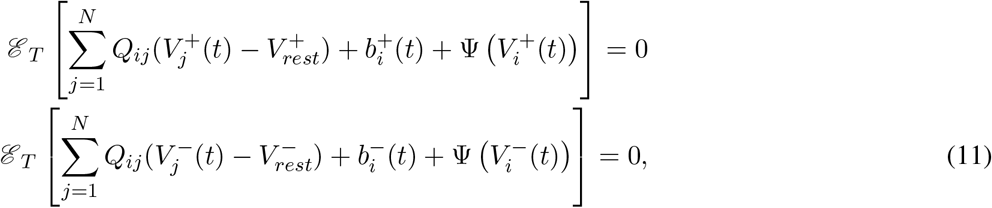

where *c* is a constant. With a transformation of variables, Eq. 11 can be written as

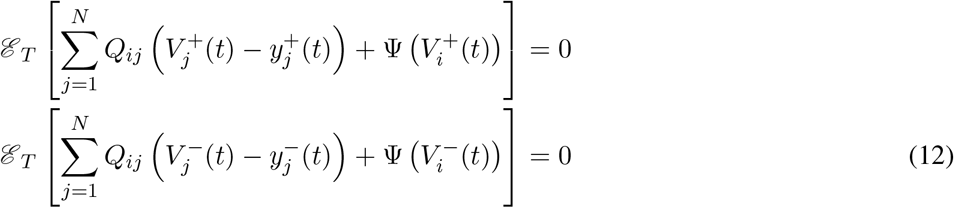

where

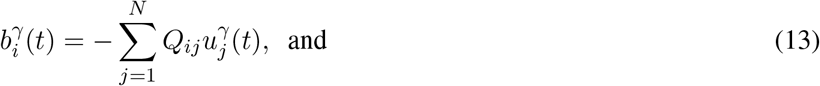

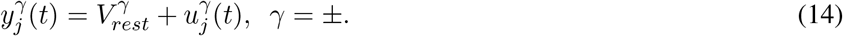

We assume *Q* to be positive definite, which ensures that 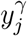 exists. Thus essentially, the external current stimulus is replaced as the weighted sum of equivalent contributions from pre-synaptic neurons. The equations in 12 can be viewed as a stochastic variant of the first-order conditions of an energy minimization problem subject to the constraints given by Eqs 7 and 8. A similar stochastic first-order framework was used in ([26]) to derive a dynamical system corresponding to ΣΔ modulation. The equivalent energy minimization problem has the following form

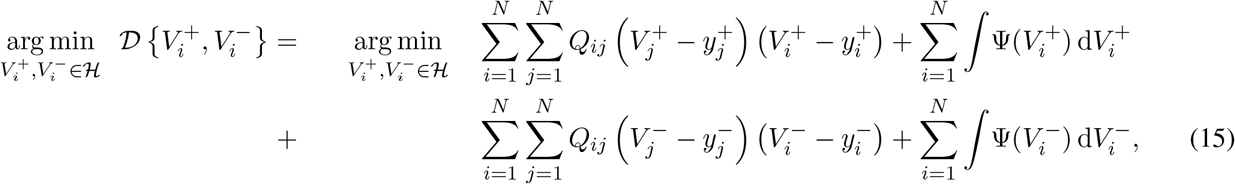

over a domain

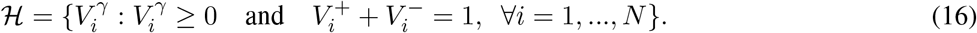

A similar system objective function was used in ([22]) to design a spiking support vector machine (SVM) for a binary classification task. Here, we explore how minimizing an energy function of this form can produce single neuron and population dynamics similar to that observed in biological neuronal networks.

The cost function 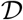 in Eq. 15 comprises two components: (a) a quadratic term that tries to minimize the error between the variables *V_i_* and *y_i_*; and (b) a regularization term Ψ(·) that produces spiking. Given any network configuration (set of synaptic connections *Q_ij_* and external inputs *b_i_*), the approach provides a way to couple the network-level neural activity to an optimization problem that strives to reach a minimum-energy state corresponding to the optimal solution when disturbed from the equilibrium. Ψ(·) is designed to be non-differentiable, with a sharp transition near the desired threshold potential (tunable) that produces large excursions in the membrane potential (i.e., spikes) from the resting state when it reaches a threshold. This is because the optimization variables 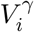 corresponding to the neurons near threshold potential cannot reach the optimal point due to the non-differentiable nature of Ψ(·), and will continue to oscillate around the threshold potential, generating spikes as long as the input stimulus is present. Ψ(·) is designed such that these oscillations take the shape of an action potential in the membrane potential *V_i_* when *V_i_* reaches the threshold.

The generation and propagation of action potentials as a way of neural signaling expends energy. The energy efficiency of biological neural networks points to the spiking dynamics being coupled with and optimized with respect to a network-level cost function. Hence at the network level, neuronal activity is essentially guided by two principles: (a) to faithfully represent the external information received in the form of spike-trains, and (b) to do so in a way that minimizes some energy cost in the dynamical system. Specifically, from Eq. 11, we see that the ion-channel activity Ψ(·) strives to continually minimize the error with which the neural responses track the time-varying input signal, so that Eq 2 is satisfied on an average. The cost function outlined in Eq. 15 also provides the framework to explore different connectivity patterns in the neural network - all within the context of optimizing a network-level cost - that has the potential to learn a better or more compact representation of the stimuli.

In ([22]), we proposed a neuron model that sequentially optimizes the cost function 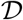 in Eq. 15 subject to constraints 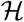 using a multiplicative update called growth transforms, a fixed-point algorithm for optimizing continuous objective functions over a manifold like 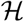 with linear and non-negativity constraints ([27]–[29]). In the following subsection, we extend this discrete-time model to a continuous-time model that incorporates axonal propagation delays.

### B. Growth Transform Neuron Model with Axonal Delay

In biology, different propagation delays along different axons produce considerable variability in the network response. These delays are intrinsically generated in other neuron models like Hodgkin-Huxley, or are synthetically introduced using a time-based kernel in leaky-integrate and fire models ([30]). In most of these cases, it is not clear how coupling the neurons together, combined with the effect of mismatch in the axonal propagation delays, would affect the stability of the network or whether the dynamics could still be interpretable. Here we extend the GT neuron model described previously to incorporate different axonal propagation delays for individual neurons in a way that keeps the neural responses tied to the system objective, and produces a stable output. The approach exploits an interpolation property of growth transforms to implement a dynamic model of a neuron by updating the variables 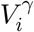, γ = ± such that the polynomial cost function 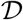 is optimized over the domain 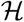. The growth transform updates ([27]) for the minimization of the energy function 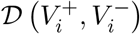 over the constrained manifold are given below

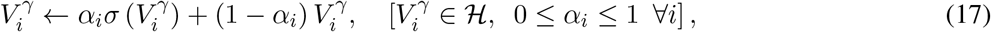

where

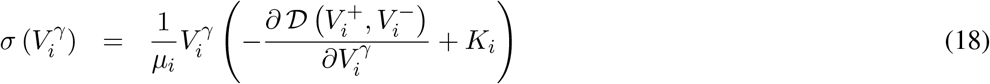

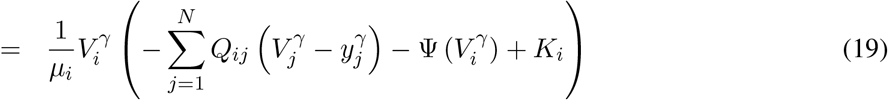

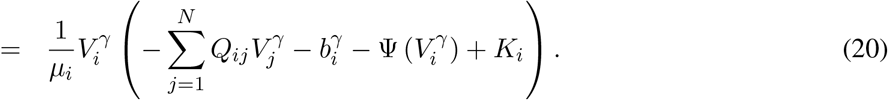

The additive factor 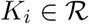 ensures the positivity constraint in Eq. 8 holds, and the normalization factor

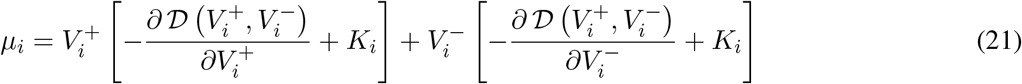

enforces the conservation constraint in Eq 7. If 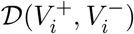 was continuously differentiable, growth transform updates would ensure that

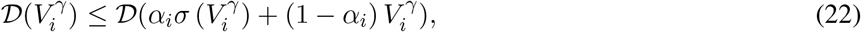

with equality condition being satisfied only when 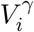 was a critical point of 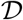. However since 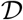 is not continuously differentiable over a sub-domain 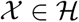, some of the variables 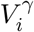 whose critical points are not directly reachable by growth transform updates will exhibit limit cycles about the sub-domain 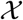. The dynamic properties of the limit cycle will be determined by the shape of Ψ(.).

Eq. 17 can be written in the following discrete-time form

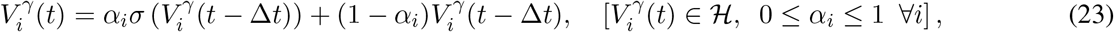

where 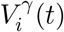 is the instantaneous value of the membrane potential at time *t*, and Δ*t* is a small time-interval. Choosing 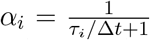, where τ = [τ_*i*_], *i* = 1, …, *N*, can be thought of as the vector of time-constants for the neuronal updates of the evolving dynamical system, we get

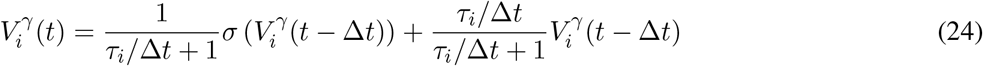

Rearranging the terms, we have

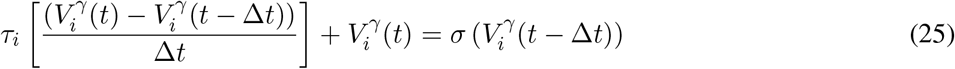

In the limiting case, as Δ*t* → 0, this reduces to the following continuous-time dynamical system

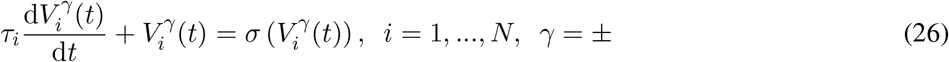

where the growth transform update 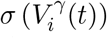 is given by Eq 20. Eq. 26 gives the equivalent growth transform neuron update for solving Eq. 2, and ensures that the network activity always converges irrespective of axonal propagation delays τ_*i*_ ∈ [0, ∞]. The neural responses are finally given by the individual membrane potentials 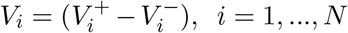. It is to be noted that axonal delays are only applied when a spike is not generated, i.e., when *V_i_* is at resting state potential or during depolarization. The update equations for the GT neuron model are summarized in Figure 3.

**Figure 3.**
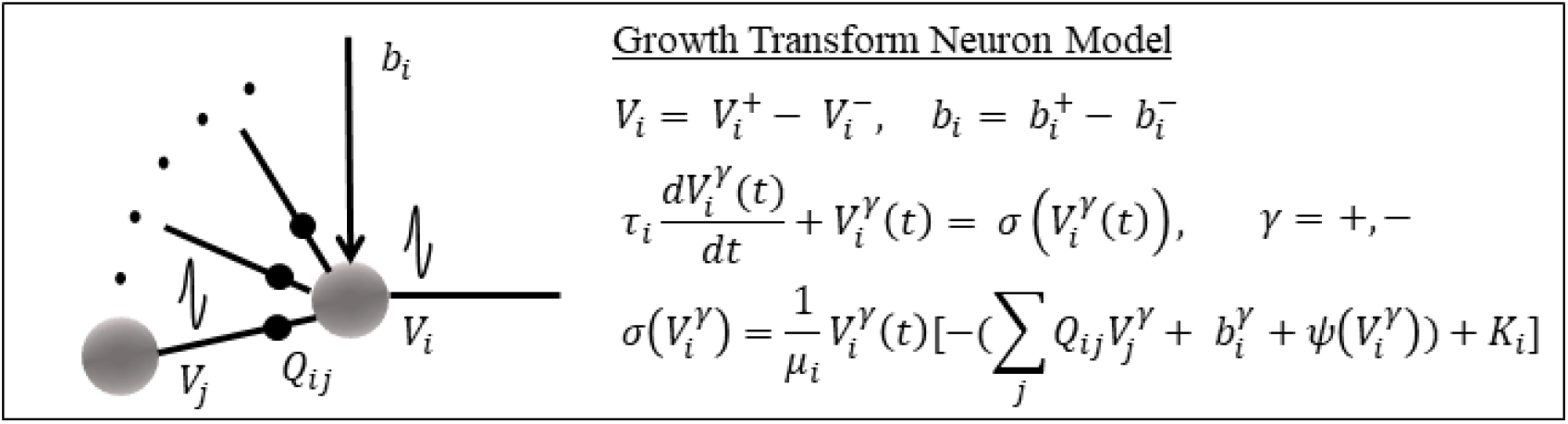
Summary of the Growth Transform Neuron model.

## III. RESULTS

In this section, we first present a detailed characterization of a single GT neuron model, and show how it can replicate many of the neuronal dynamics observed in literature [31] by altering various aspects of the energy functional. We then demonstrate how the GT neuron encodes a time-varying stimulus in its spike pattern using different encoding techniques. Finally, we use a network of GT neurons to perform auditory feature extraction for a speaker recognition task. We provide extensive simulation results with an uncoupled network, and coupled networks with only excitatory, only inhibitory and with either types of connections between neurons, both for no axonal delays and with random axonal propagation delays.

### A. Single GT Neuron Characterization

The ion-channel function Ψ(.) used in this paper is shown in Figure 4. The gradient discontinuity in this case was set to *V*^+^ = *V*^−^ = 0.5. In resting state, 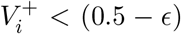 and 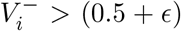, where ϵ is the width of the very narrow transition zone that sets the minimum input current needed for a spike as well as its height. When a depolarizing current pulse comes, both 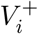 and 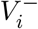 try to move towards 0.5. If the input current is strong enough, the variables cross over the threshold, producing a spike. This adds a large penalty to the gradient term for 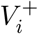, and the updates force the variables to come back to a hyperpolarized state (a more negative *V_i_*) than where it started from. The resulting membrane potential trace *V_i_* is also given in Figure 4.

**Figure 4.**
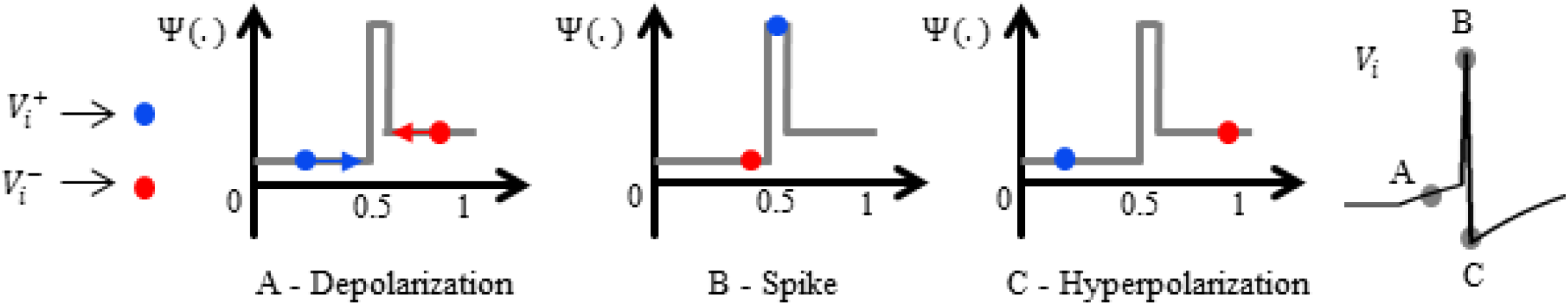
Ion-channel function Ψ(.) used in this paper, with the paths traced by optimization variables 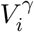 in producing a spike.

We begin by characterizing the proposed model using a time-varying control input as shown in Figure 5A. The input is processed by a GT neuron, whose response *V_i_* and the corresponding spike raster are given in Figure 5B. Figure 5C shows the expectation of the net instantaneous current at the neuron given by Eq. 2, taken over a time-window. Although the instantaneous values of the total current at the neuron can be fluctuating considerably depending on the stimulus inputs and ionic currents, growth-transform updates ensure the time-average is always close to zero.

**Figure 5.**
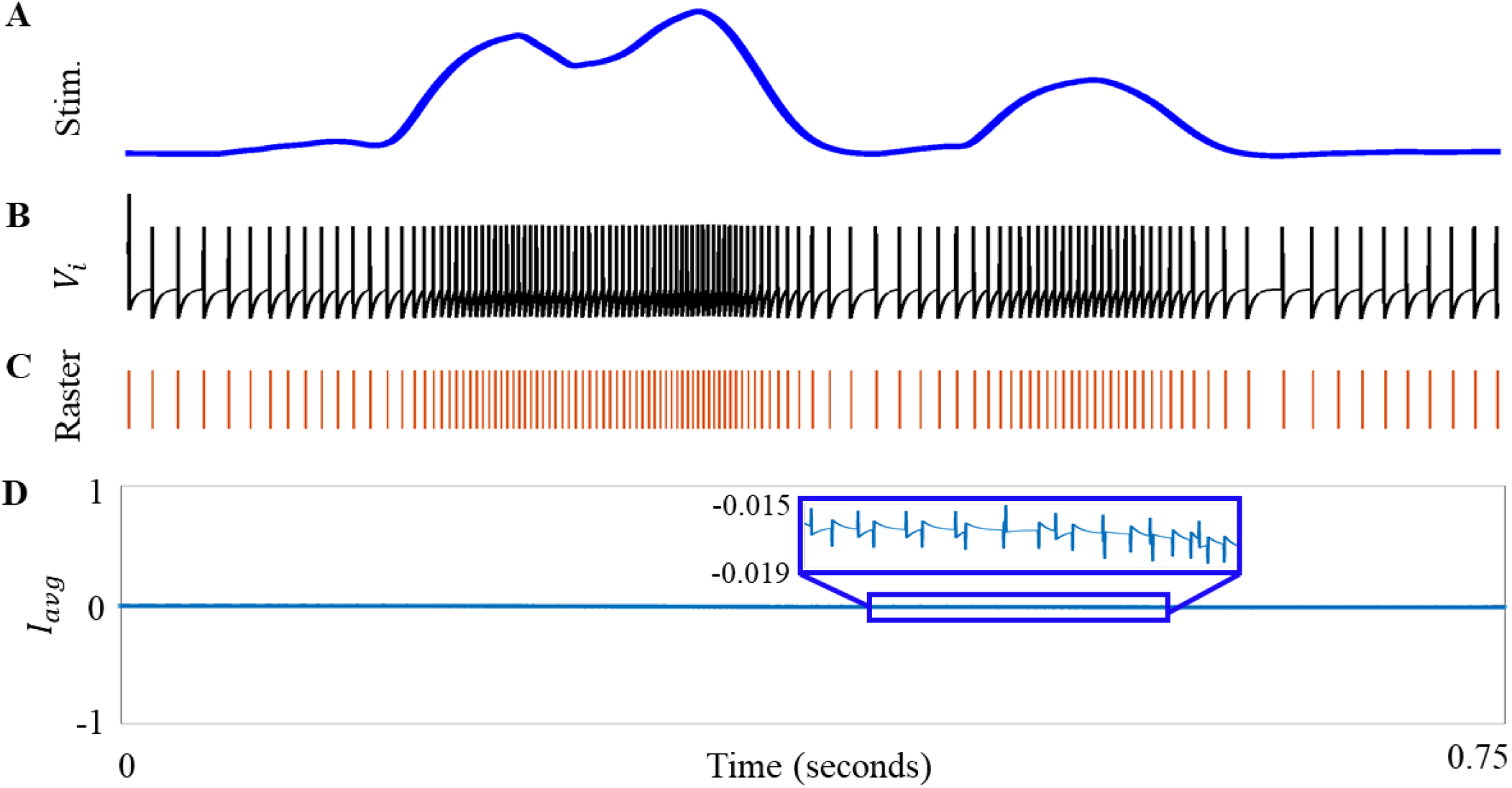
(A) Time-varying stimulus. (B) Membrane potential trace and spike raster. (C) Time-average of the net current at the neuron, given by Eq. 2.

We next investigate the effect of axonal propagation delays at a particular neuron. Figure 6A shows the modulation in the spiking frequency of a neuron due to increasing axonal delay (i.e., increasing τ) in response to the same input.

**Figure 6.**
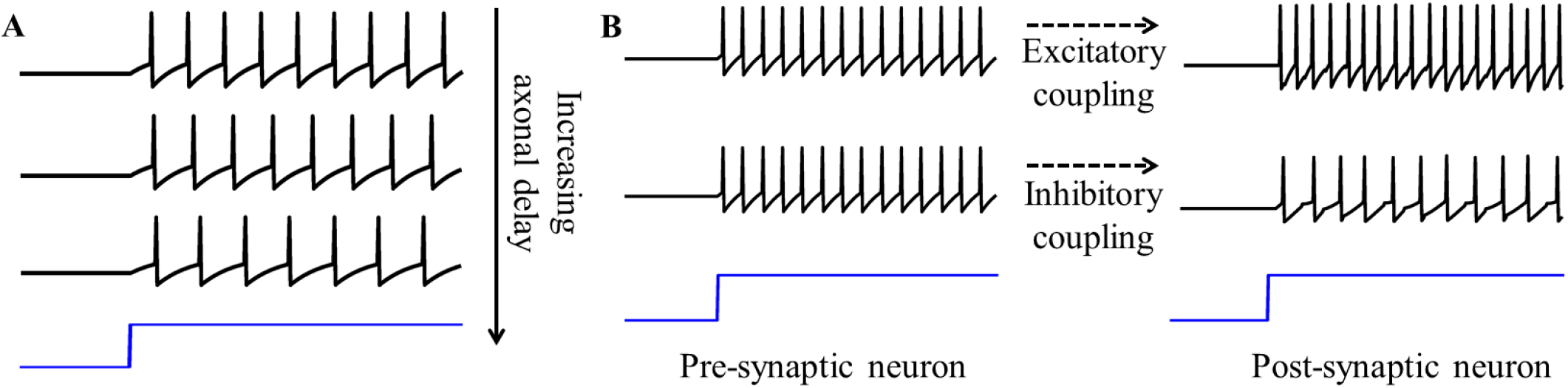
(A) Effect of axonal delay on spike propagation. (B) Effect of excitatory and inhibitory couplings on the post-synaptic neuron.

The GT neural network provides a way to couple a network of neurons together that enables the pre-synaptic neuron to excite/inhibit the post-synaptic neuron, depending on the sign of the synaptic weight connecting them. This is illustrated in Figure 6B. An excitatory (positive) coupling increases the firing rate of the post-synaptic neuron, whereas an inhibitory (negative) coupling reduces its firing rate.

### B. Single GT neuron Dynamics

Single neurons show a vast repertoire of response characteristics and dynamical properties that lend richness to their computational properties at the network level. Izhikevich in ([31]) provided an extensive review of different spiking neuron models and their ability to produce the different dynamics observed in biology. We now show how we can reproduce a number of such dynamics by changing different aspects of the cost function in Eq. 15.

- **Tonic spiking.** When stimulated with a constant DC current pulse, a vast majority of neurons fire single, repetitive action potentials for the duration of the stimulus, with or without adaptation. Cortical neurons like regular spiking (RS) excitatory neurons, as well as low-threshold spiking (LTS) and fast spiking (FS) inhibitory interneurons show tonic spiking behavior ([32]–[34]). A simulation of such a firing pattern obtained using our model is shown in Figure 7A.
- **Tonal excitation.** Response of a GT neuron to a sinusoidally varying stimulus is shown in Figure 7B.
- **Class 1 excitability.** Hodgkin identified three classes of neurons on the basis of their patterns of repetitive spiking by studying the response of squid axons to current pulses of various amplitudes ([35], [36]). Class 1 neurons are able to generate action potentials with arbitrarily low frequencies in response to weak but superthreshold stimulus. The frequency of firing varies smoothly with the strength of injected current giving rise to a continuous *f - I* curve. GT neuron inherently shows Class 1 excitability, which is demonstrated by its response to a ramp input in Figure 7C.
- **Integration.** Fig. 7D shows how a single GT neuron works as a leaky integrator, preferentially spiking to high-frequency or closely-spaced input pulses.
- **Bursting.** Bursting neurons fire discrete groups of spikes interspersed with periods of silence in response to a constant stimulus ([32], [33], [37], [38]). Bursting arises from an interplay of fast ionic currents responsible for spiking and slower intrinsic membrane currents that modulate spiking activity, causing the neuron to alternate between activity and quiescence. We model the intrinsic ionic mechanism of the faster process of action potential generation, and the slower process governing burst occurrence and duration by slowly modulating the shape of the underlying ion-channel function Ψ(.) in a periodic manner in presence of excitation, so that the cell alternates between spiking and silence, thus encoding the input in bursts instead of individual spikes. Simulation of a bursting neuron in response to a constant step input obtained using the growth transform network is shown in Fig. 7E.
- **Spike frequency adaptation.** When presented with a prolonged stimulus of constant amplitude, many cortical cells including regular spiking (RS) neurons and low-threshold spiking (LTS) inhibitory neurons initially respond with a high-frequency spiking that decays to a lower steady-state frequency ([39]). This adaptation in the firing rate is caused by a negative feedback to the cell excitability due to gradual inactivation of depolarizing currents or activation of slow hyperpolarizing currents upon depolarization, and occur at a time-scale slower than the rate of action potential generation. This can be replicated in our model by tuning a single parameter, *C*. In the final cost function, this factor *C* that models the single-cell excitability, appears as the regularization hyperparameter, setting the relative weightage of the two terms, the net excitation received by a neuron and the net ionic activity in response. In order to model spike frequency adaptation, the excitability factor itself is made to undergo a first-order dynamics as follows

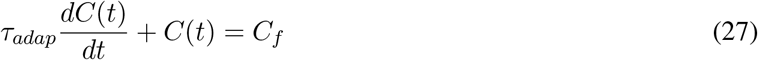

where τ_*adap*_ >> τ. The corresponding response is given in Figure 7F.

**Figure 7.**
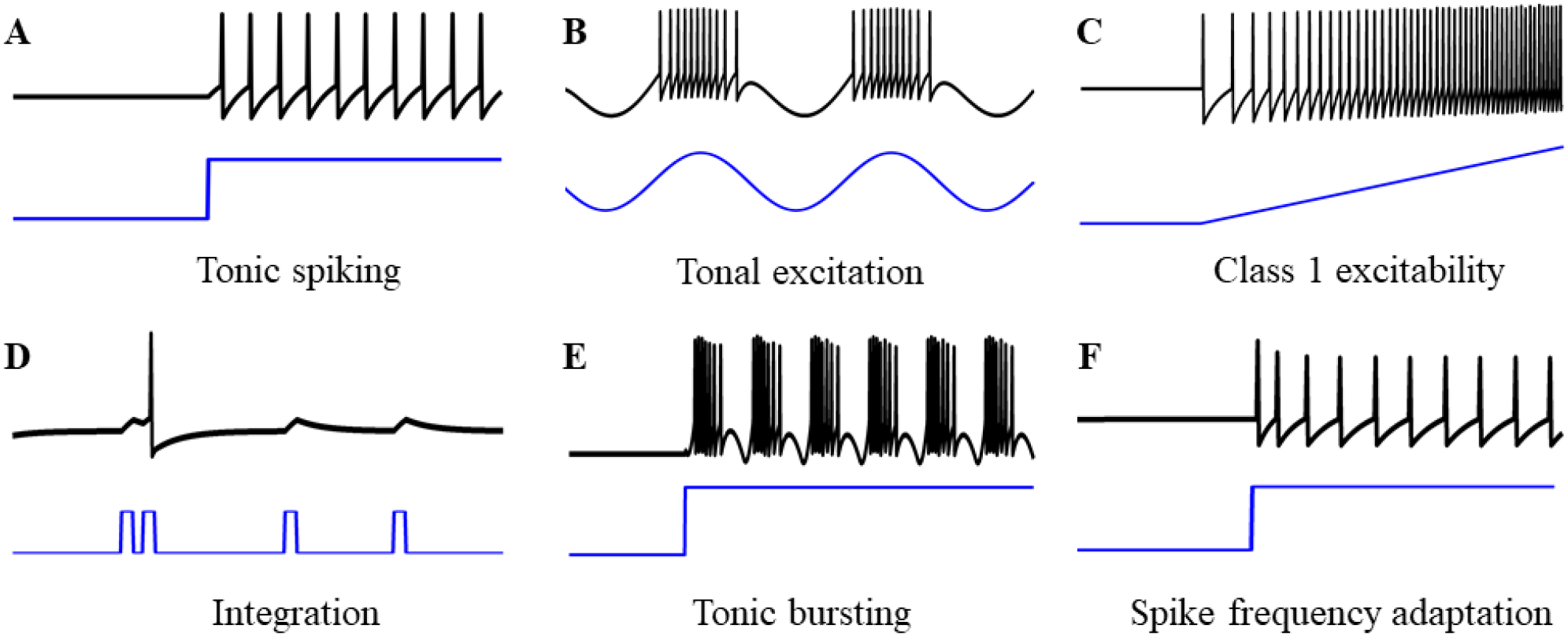
Single-neuron responses. Simulations showing different dynamical properties of the GT neuron.

Software (MATLAB) implementation of single and coupled GT neuron models are available through the webportal http://aimlab.seas.wustl.edu/document/gtn_gui_matlab.zip. The tool enables users to visually see the effect of different hyper-parameters on different GT neuron dynamics and on different population dynamics produced by a network of coupled GT neurons.

### C. Spike-encoding using Growth Transform Neurons

Before we use a network of GT neurons to encode auditory features, we briefly discuss different spike-encoding patterns produced by a single GT neuron in response to the control stimulus given previously. For a time-varying input stimulus, GT neurons show a number of encoding properties similar to those observed in biological neural networks.

- **Rate encoding**: Mean firing rate over time is a popular rate coding scheme that claims that the spiking frequency or rate is modulated with the stimulus intensity ([40]). Rate-based features were obtained from this spike-train by computing the average spiking frequency of the neuron over a moving window.
- **Inter-spike interval-based encoding**: Temporal codes like inter-spike intervals claim that timings between successive spikes carry valuable information about the stimuli ([41]). We explored whether actual spike-timings produced by GT neurons can garner useful information about the input signal. For this, we computed the average inter-spike interval (ISI) for a neuron over a moving window over the entire spike-train.
- **Mean membrane potential**: For the next set of features, we explored whether inclusion of subthreshold activity improves stimulus-encoding. For this, a moving average was computed over the thresholded membrane potential V itself, that included both spikes and subthreshold activity. The threshold value was chosen to be slightly lower than the threshold membrane potential.

### D. Auditory Feature Extraction

The feature extraction algorithm proposed in this paper is summarized in Figure 8. Gammatone filterbanks ([42]) simulating cochlear filtering in the auditory system produce outputs that track the energy in different frequency bands over the entire bandwidth. Each filterbank output *x_i_* is rectified, low-pass filtered, scaled to [0, 1] and processed by a GT neuron in a differential manner as *b_i_, i* = 1, …, *N*, N being the number of filterbanks used. The growth transform neurons track the energy in individual filterbank outputs, thus performing a basic decoding of speech signals in terms of frequency, intensity and duration. The proposed network outlines an approach where the early stages of acoustic information processing, which is analogous to the functionality of cochlear nuclei, can be connected to a network-level objective function, and the spiking dynamics produced is the direct derivative of a continuous optimization of the network cost as it processes a continually varying input signal in real-time.

**Figure 8.**
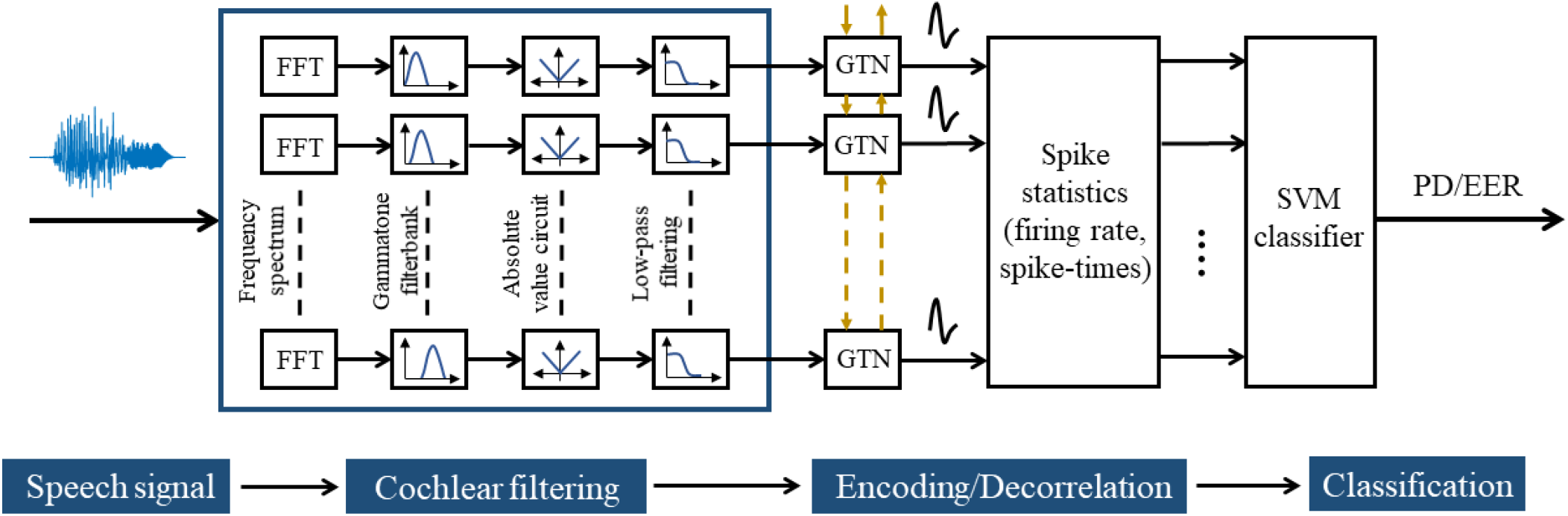
Signal-flow showing the feature extraction procedure for a speaker recognition task. The coupled growth transform neural network processes rectified and low-pass filtered gammatone filterbank outputs and produces spike trains from which different spike statistics are extracted to encode discriminatory features.

In the next two subsections, we explore how spike-encoded features can be extracted from speech signals with uncoupled and coupled GT networks. For this, we use a sample speech signal corresponding to the utterance “one”. Figures 9A and B show the speech signal and its corresponding spectrogram.

**Figure 9.**
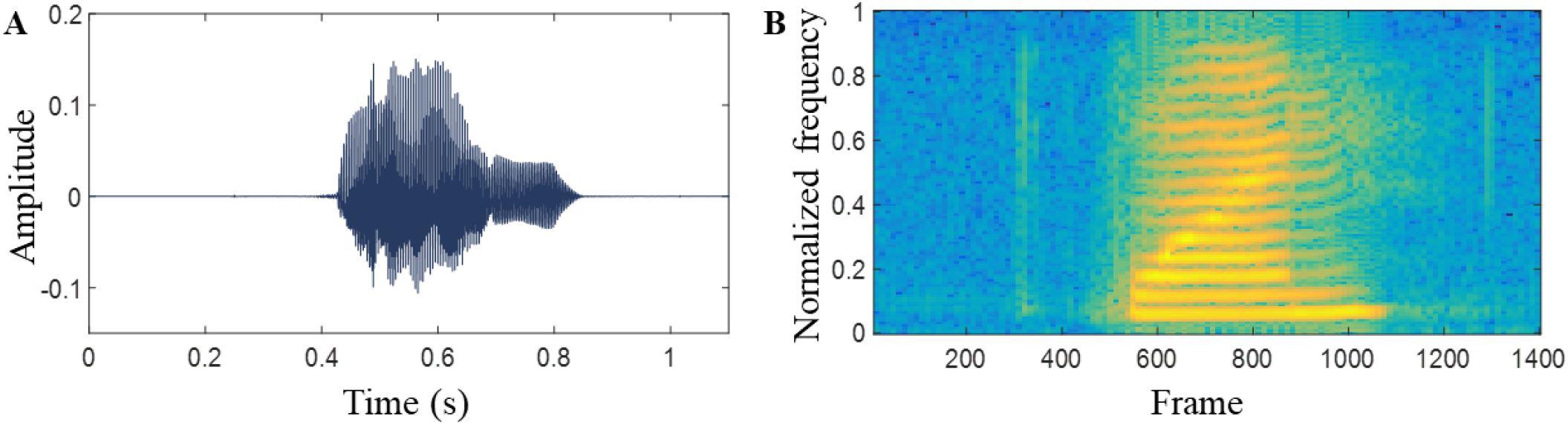
(A) A typical example of a speech signal corresponding to the utterance “one”. (B) The corresponding spectrogram with frame size = 150.

1. *Uncoupled GT Neural Network Features*: We first consider an uncoupled network, where the synaptic weight matrix Q is diagonal. Each neuron therefore independently processes each filterbank output without contributions from the other neurons. For the speech signal in Figure 9A, the spike raster produced by a GT neural network with 20 neurons, each processing a single output channel from a filterbank of size 20, is presented in Figure 10A. Figures 10B, C and D show colormaps for the three feature sets.
2. *Coupled GT Neural Network Features*: Next, we explore how different network connectivity patterns affect the network representation of input stimuli. For this, we considered three coupled networks with excitatory connections, inhibitory connections, and with both excitatory and inhibitory connections. For the last network, neurons were randomly chosen to be either excitatory or inhibitory in nature. The corresponding spike rasters and feature colormaps are shown in Figures 11, 12 and 13 respectively. We see that different types of coupling affect the spike-patterns and encoding in markedly different ways. As expected, the average firing rate for a coupled network with inhibitory connections is much lower than the uncoupled network.

**Figure 10.**
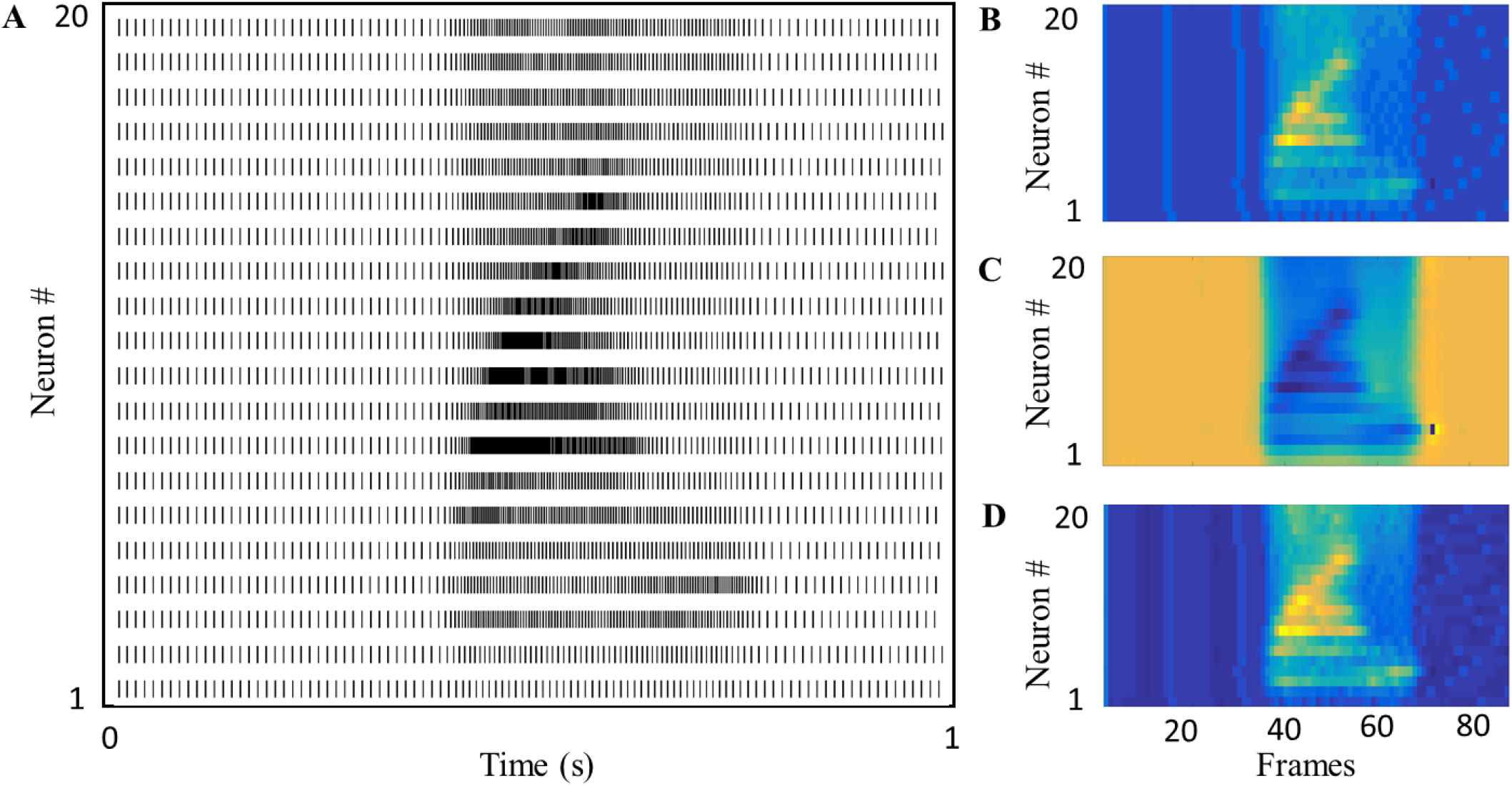
(A) Spike raster produced by the uncoupled growth transform neural network. (B), (C) and (D) Colormaps depicting feature sets with rate, spike-timing and mean membrane potential-based encoding of the speech signal in 9(A) respectively.

**Figure 11.**
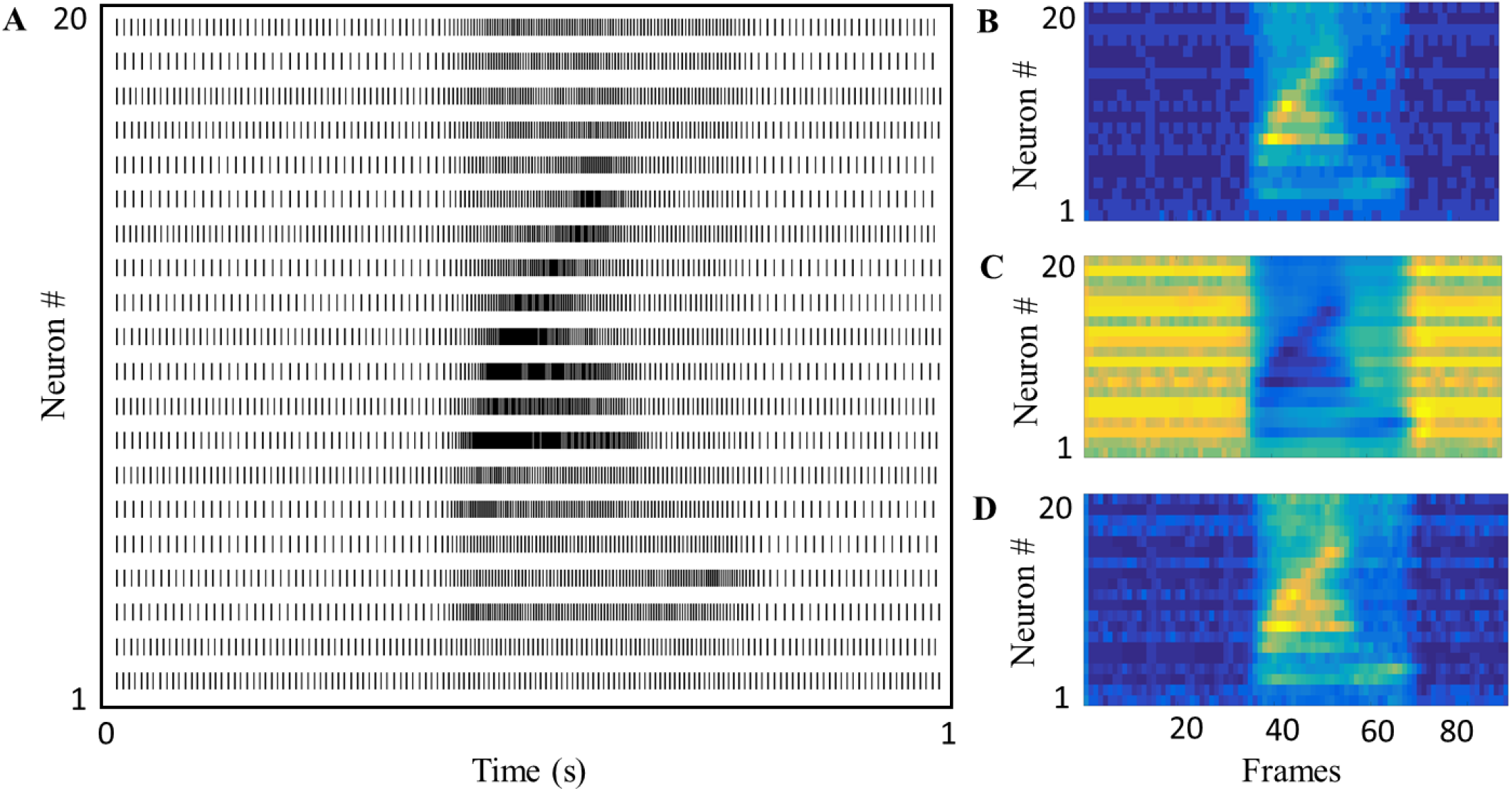
(A) Spike raster produced by the coupled growth transform neural network with sparse random excitatory connections. (B), (C) and (D) Colormaps depicting feature sets with rate, spike-timing and mean membrane potential-based encoding of the speech signal in 9(A) respectively.

**Figure 12.**
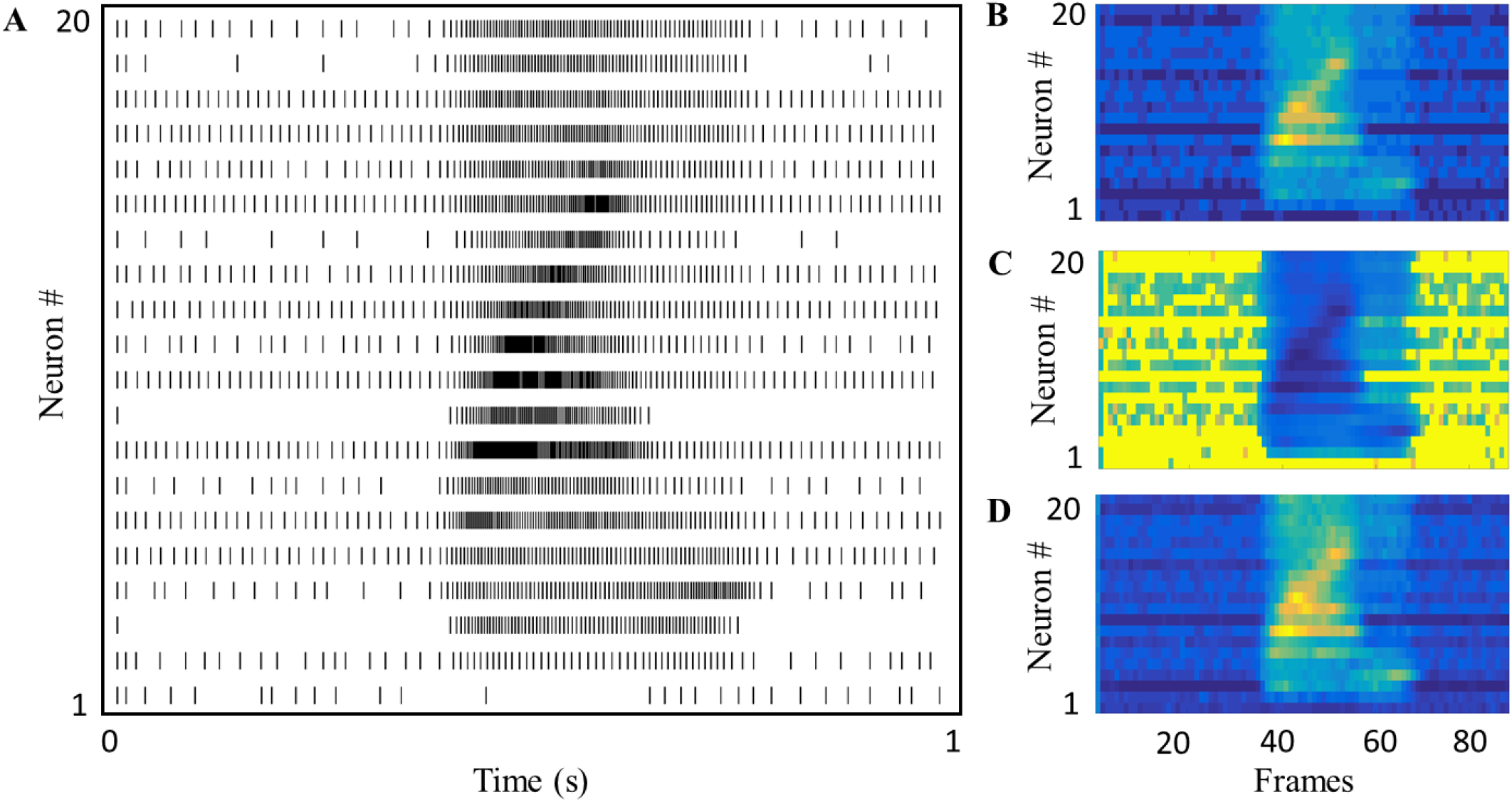
(A) Spike raster produced by the coupled growth transform neural network with sparse random inhibitory connections. (B), (C) and (D) Colormaps depicting feature sets with rate, spike-timing and mean membrane potential-based encoding of the speech signal in 9(A) respectively.

**Figure 13.**
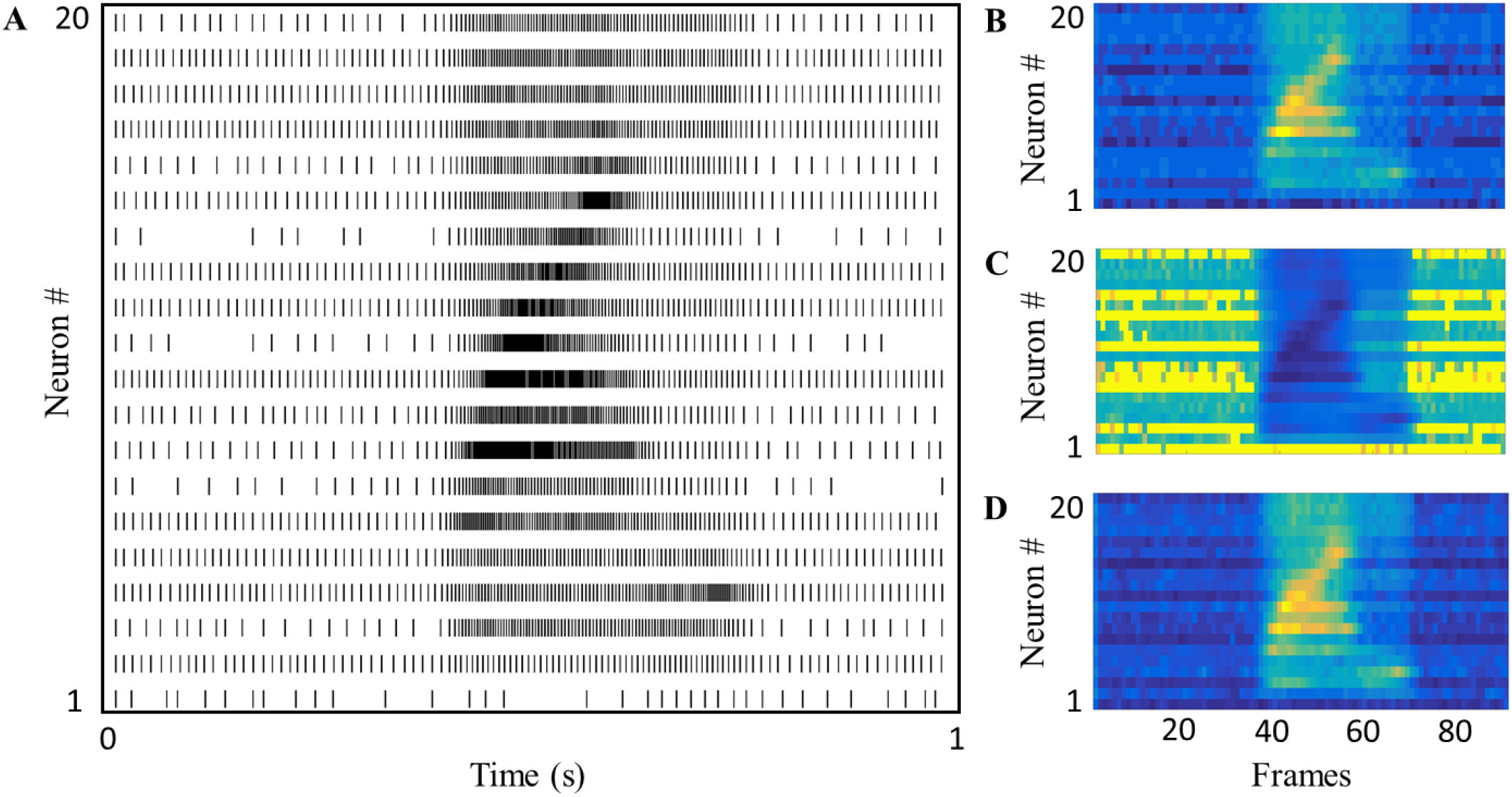
(A) Spike raster produced by the coupled growth transform neural network with sparse random excitatory as well as inhibitory connections. (B), (C) and (D) Colormaps depicting feature sets with rate, spike-timing and mean membrane potential-based encoding of the speech signal in 9(A) respectively.

### E. Speaker Recognition Experiments

A speaker identification task utilizing the growth transform feature extraction algorithm is evaluated against the YOHO database, available through the Linguistic Data Consortium ([43]). This corpus is a collection of combination lock phrases (e.g. 26 – 81 – 56, pronounced twenty-six eighty-one fifty-six) for 138 subjects collected across a three month period. The data of interest are limited to the four enrollment sessions, with 24 utterances per session, collected in an office environment at an 8 kHz sampling rate with 3.8 kHz analog bandwidth. The first session is used for training, the second session for cross-validation, and the remaining third and fourth sessions were applied as a test set to evaluate the performance of the speaker identification system. There is a nominal time interval of three days between the sessions in YOHO. The data of each utterance in a session is initially normalized to the maximum response in the utterance before being passed through a second order gammatone filterbank with a quality factor of seven. Center frequencies of the filters in the filterbank are equally spaced according to the mel-scale up to 4 kHz, where the filterbank size is 20 for all the experiments reported in this paper. The envelopes of the rectified filterbank outputs are fed into the growth transform neural network as external stimuli. The spike-trains produced by the network are smoothed out using a moving window average and used as inputs to a *Gini* support vector machine (Gini-SVM) ([44]) which does the task of identification. The performance metrics reported in this work are the average across all considered speakers' probability of detection (PD) and the equal error rate (EER), which is the point at which the false rejection and false acceptance rates are equivalent. In general, a higher PD or lower EER would be indicative of better performance.

First, the feature set was evaluated using a baseline recognition system without any spiking neurons. The purpose of this evaluation was to determine the upper-limit of recognition that can be achieved by the gammatone filterbank auditory front-end. Then, each feature set was tested for four cases: (a) an uncoupled network, (b) a coupled network with sparse random inhibitory connections, (c) a coupled network with sparse random excitatory connections, and (d) a coupled network where neurons were randomly chosen to be either excitatory or inhibitory with random synaptic weights. For both uncoupled and coupled networks, DCT was applied as a post-processing step to decorrelate the features, and the DC component was dropped to get 19 usable filter outputs. Δ features (i.e. the first derivative or velocity) were concatenated to the limit-cycle statistics extracted from the spike-trains. To reduce the effort required by the Gini-SVM, only a subsample of the feature vector (taken at uniform intervals) was fed into the system, for the YOHO utterances that last a couple of seconds, the resulting feature vector length can be several tens of thousands, while the subsampled version might be only a couple hundred.

Results for all the experiments over the entire database are given in Table I. We see that while the performance of the uncoupled network is similar to the baseline, introducing random couplings between neurons do not degrade the classification performance, especially when inter-spike intervals are used as features. This enables us to explore network connectivities that might lead to a more energy-efficient representation of external stimuli. For example, the coupled network with only inhibitory connections, is seen to reduce the overall firing rate of the network by an average of 5% per utterance, while still being able to extract relevant features for encoding of auditory data. Moreover, inclusion of subthreshold activity is seen to result in a finer discrimination in comparison to rate-encoding in almost all the cases.

**Table I.**
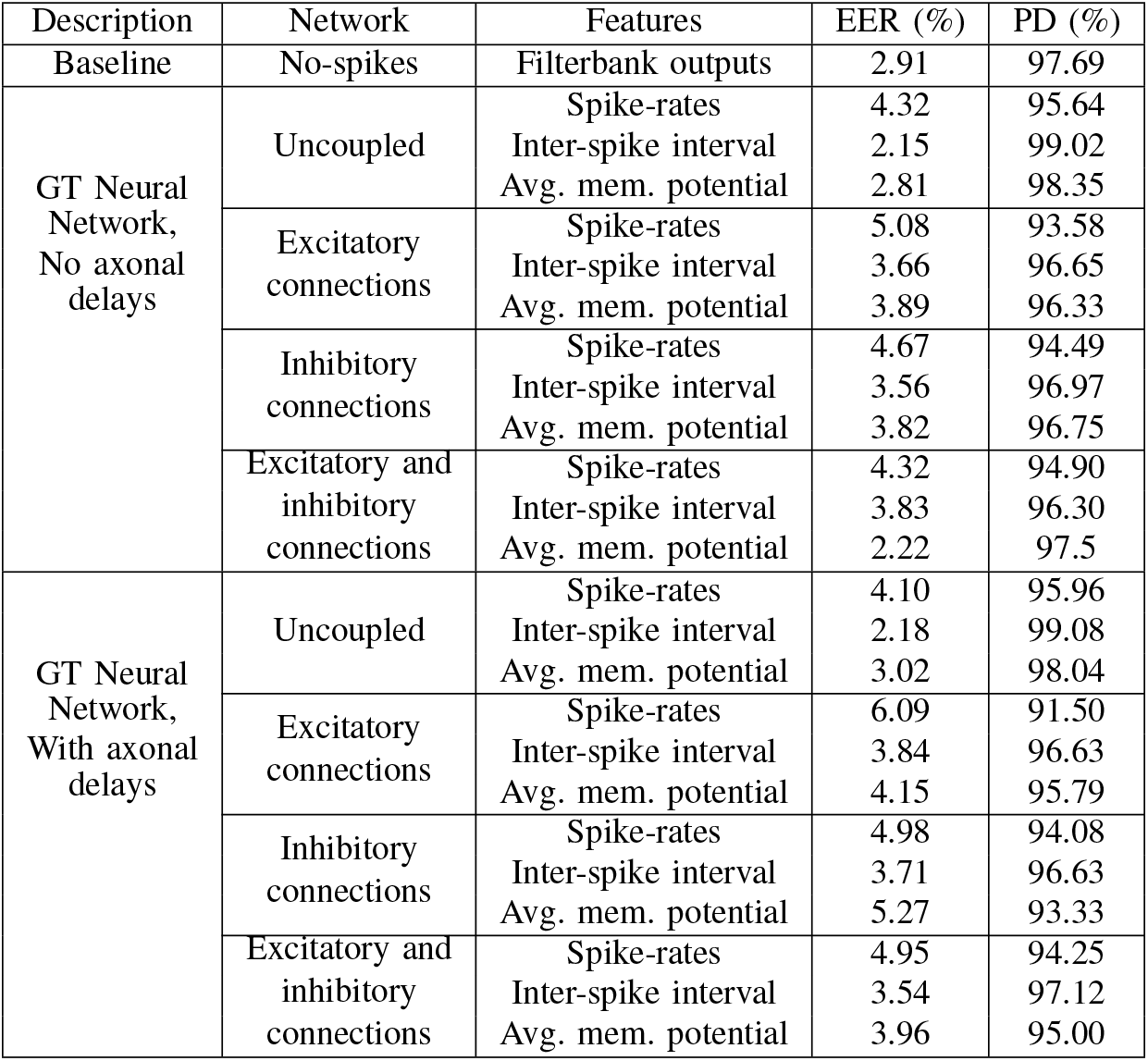
Speaker recognition results on the YOHO database using GT neural network.

| Description | Network | Features | EER (%) | PD (%) |
| --- | --- | --- | --- | --- |
| Baseline | No-spikes | Filterbank outputs | 2.91 | 97.69 |
| GT Neural Network,<br>No axonal delays | Uncoupled | Spike-rates | 4.32 | 95.64 |
|  |  | Inter-spike interval | 2.15 | 99.02 |
|  |  | Avg. mem. potential | 2.81 | 98.35 |
|  | Excitatory connections | Spike-rates | 5.08 | 93.58 |
|  |  | Inter-spike interval | 3.66 | 96.65 |
|  |  | Avg. mem. potential | 3.89 | 96.33 |
|  | Inhibitory connections | Spike-rates | 4.67 | 94.49 |
|  |  | Inter-spike interval | 3.56 | 96.97 |
|  |  | Avg. mem. potential | 3.82 | 96.75 |
|  | Excitatory and inhibitory connections | Spike-rates | 4.32 | 94.90 |
|  |  | Inter-spike interval | 3.83 | 96.30 |
|  |  | Avg. mem. potential | 2.22 | 97.5 |
| GT Neural Network,<br>With axonal delays | Uncoupled | Spike-rates | 4.10 | 95.96 |
|  |  | Inter-spike interval | 2.18 | 99.08 |
|  |  | Avg. mem. potential | 3.02 | 98.04 |
|  | Excitatory connections | Spike-rates | 6.09 | 91.50 |
|  |  | Inter-spike interval | 3.84 | 96.63 |
|  |  | Avg. mem. potential | 4.15 | 95.79 |
|  | Inhibitory connections | Spike-rates | 4.98 | 94.08 |
|  |  | Inter-spike interval | 3.71 | 96.63 |
|  |  | Avg. mem. potential | 5.27 | 93.33 |
|  | Excitatory and inhibitory connections | Spike-rates | 4.95 | 94.25 |
|  |  | Inter-spike interval | 3.54 | 97.12 |
|  |  | Avg. mem. potential | 3.96 | 95.00 |

In order to test how the GT neural network performs with random axonal propagation delays, we conducted the same set of experiments, but with different τ*_i_* = 1, …, *N* chosen randomly for each neuron. The rate of sampling of the input signal was the same as the no-delay case. We see that even with random axonal delays, the classification accuracy using the spike-encoded features does not degrade, even though there is a mismatch between the rates at which different neurons encode the auditory information.

## IV. DISCUSSION

This paper attempts to answer the following fundamental question - Can the spiking dynamics in a synthetic neuron model be derived from a network-level energy-minimization problem? In other words, can we design an appropriate energy functional, which when optimized under realistic physical constraints, produces known neural dynamics? For this, we propose a new spiking neuron model based on growth transform updates where the process of spike generation and encoding is derived from a network cost function that strives to achieve a balance between the net excitation received by a neuron and the net ionic processes that determine its response, so that the time-average of accumulated charge in the cell membrane is zero. We derive a dynamical system formulation of the growth transform updates which takes into account axonal propagation delays in individual neurons. We analyze the model for different inputs and parameter variations, and demonstrate that it is well able to fulfill the current-balance equation from which we derive the neuronal updates.

Electrophysiological recordings show a rich repertoire of response dynamics at the cellular level. We showed how the GT neuron can reproduce a number of such dynamics by altering various parameters of the cost function, and tried to derive mechanistic insights about the significance of each. It was also demonstrated that the GT neuron can encode external stimuli using firing rates and inter-spike intervals, similar to biological neurons. Moreover, the sub-threshold activity of GT neurons was also seen to encode stimulus information in addition to spike-based statistics. These encoding techniques were applied for the task of encoding auditory features in a speaker recognition problem. The neural activity of GT neurons is always coupled to a network-level cost function, which enables us to explore different synaptic connectivities coupling the neurons, without sacrificing network stability. For example, introducing inhibitory couplings between neurons was seen to reduce the overall firing rate of the network, without degrading classification accuracy.

The paper illustrates how spike generation and spike-based encoding in an artificial neuron model could take place within a network-level optimization framework, thereby providing a tool to develop scalable neuromorphic machine learning algorithms. We have only considered fixed synaptic weights between neurons in this paper. An interesting direction of future research would be to learn these weights for an ‘optimal' representation of the external stimuli, where the optimality condition could be based on spike counts, number of connections, etc. Finally, an end-to-end synthetic auditory system could be implemented within the same framework by replacing the classification module by a network of GT neurons capable of performing higher processing functions like speech or speaker recognition through spike-based encoding and learning.

## CONFLICT OF INTEREST STATEMENT

The authors declare that the research was conducted in the absence of any commercial or financial relationships that could be construed as a potential conflict of interest.

## AUTHOR CONTRIBUTIONS

AG and SC contributed to the conception and design of the study; KA, AG and DM conducted the simulations; DM designed the MATLAB interface, AG and SC wrote the first draft of the manuscript; AG, KA and SC wrote sections of the manuscript. All authors contributed to manuscript revision, read and approved the submitted version.

## FUNDING

This material is based upon work supported by the National Science Foundation under Grant Nos. CSR-1405273, DGE-0802267, DGE-1143954. Any opinions, findings, and conclusions or recommendations expressed in this material are those of the author(s) and do not necessarily reflect the views of the National Science Foundation.

